# Motor preparation tracks decision boundary crossing in temporal decision-making

**DOI:** 10.1101/2024.09.24.614731

**Authors:** Nir Ofir, Ayelet N. Landau

## Abstract

Interval timing, the ability of animals to estimate the passage of time, is thought to involve diverse neural processes rather than a single central “clock” (Paton & Buonomano, 2018). Each of the different processes engaged in interval timing follows a different dynamic path, according to its specific function. For example, attention tracks anticipated events, such as offsets of intervals (Rohenkohl & Nobre, 2011), while motor processes control the timing of the behavioral output (De Lafuente et al., 2024). Hence, different processes provide complimentary perspectives on mechanisms of time perception. The temporal bisection task, where participants categorize intervals as “long” or “short”, is thought to rely on the same mechanisms as other perceptual decisions (Balcı & Simen, 2014). In line with this hypothesis, we previously described an EEG potential that tracks the formation of decision following the end of the timed interval (Ofir & Landau, 2022). Here, we track the dynamics of motor preparation to investigate the formation of decision within the timed interval. In contrast to typical perceptual decisions, where motor plans for all response alternatives are prepared simultaneously (Shadlen & Kiani, 2013), we find that different temporal decisions develop sequentially. While preparation for “long” responses was already underway before interval offset, no preparation was found for “short” responses. Furthermore, within intervals categorized as “long”, motor preparation was stronger at interval offset for faster responses. Our findings shed light on the unique dynamics of temporal decisions and demonstrate the importance of considering neural activity in timing tasks from multiple perspectives.

## Introduction

Tracking the passage of time is a fundamental capability of animals that forms a scaffold upon which behavior is organized. An “internal clock” underlying all timed behaviors has yet to be described, and a growing body of work suggests such a “clock” might not exist (Paton & Buonomano, 2018). However, any single timing behavior involves multiple processes, spanning from early perceptual stages to the final motor response, and the perception of time can leave measurable traces in each. For example, the temporal bisection task, in which participants categorize intervals as being “long” or “short”, recruits sensory processes to identify the onset and offset of the interval and motor processes to report the decision. Previously, we have reported an EEG potential which builds up from interval offset to the participants’ response (Ofir & Landau, 2022). The potential was evoked by intervals that were categorized as “short” but not by those categorized as “long”. In addition, for “short” intervals, the potential was larger in amplitude for shorter intervals, while for “long” intervals there was no dependence on interval duration. This pattern is consistent with theoretical studies describing temporal bisection as a bounded accumulation process (Balcı & Simen, 2014). In its simplest form, the bounded accumulation assumes a noisy accumulator and a decision boundary. The noisy accumulator, representing elapsed duration, starts at interval onset. If the boundary is reached during the interval, the interval is categorized as “long”, and as “short” otherwise. We found that that the offset-evoked potential reflected the distance from decision boundary at interval offset (Ofir & Landau, 2022). However, this potential only provides an indirect, or post-hoc, measure of the actual accumulation of temporal evidence since it starts at interval offset. In the present study we would like to directly investigate the process of temporal evidence accumulation. To this end, we tracked the preparation of the manual response as reflected in activity in the mu-beta band (8-30 Hz; Pfurtscheller & Lopes Da Silva, 1999).

Mu-beta (8-30 Hz) amplitude is a robust signature of preparing and executing a motor command (Pfurtscheller & Lopes Da Silva, 1999). Mu-beta amplitude in the hemisphere contralateral to the responding hand decreases gradually towards a fixed level at which movement onsets. When different decisions are mapped to different hands, the amplitude in each hemisphere decreases independently, and the dynamics of both hemispheres resemble that of a race: The first hemisphere to reach the threshold is the one to respond (O’Connell & Kelly, 2021). Therefore, mu-beta lateralization (i.e. the relative amplitude of mu-beta between the two hemispheres) is taken to reflect differential preparation (and accordingly differential evidence) towards one response alternative vs. the other (Donner et al., 2009).

Based on the literature, three possible scenarios for the relative preparation of “short” and “long” responses during the timed interval can be expected: The first is that, initially, there will be stronger preparation to a “short” response, which will gradually switch into stronger preparation to a “long” response. Preparation for “short” can either be apparent already at baseline, or it could develop rapidly after interval onset (Dashed and solid lines respectively in **Figure 1G**). Support for this prediction can be found in neuronal recordings from lateral intraparietal neurons in rhesus monkeys performing a similar task (Leon & Shadlen, 2003). Alternatively, theoretical accounts based on response time analysis suggest that during the interval itself only the “long” alternative is entertained (Balcı & Simen, 2014). In perceptual decision-making tasks, motor preparation has been found to reflect the gradual accumulation of evidence towards a choice (Afacan-Seref et al., 2018; Balsdon et al., 2023; Steinemann et al., 2018; Wilming et al., 2020; Wyart et al., 2012), implying that only preparation for “long” responses will be evident during the interval. Based on the suggestion that time perception corresponds tightly to perceptual decision-making (Balci & Simen, 2016), the second scenario predicts that lateralization will immediately start growing towards the “long” hand following interval onset (**Figure 1H**). Finally, it could be that motor preparation in temporal decisions is delayed until a decision has been made, rather than track accumulated evidence, in contrast to other perceptual decisions. Hence, in the third scenario we would expect to see preparation for “long” responses emerge later during the interval, once “long” decisions are committed (**Figure 1I**). In the second and third scenarios, no preparation towards the “short” hand is expected during the interval.

**Figure 1:**
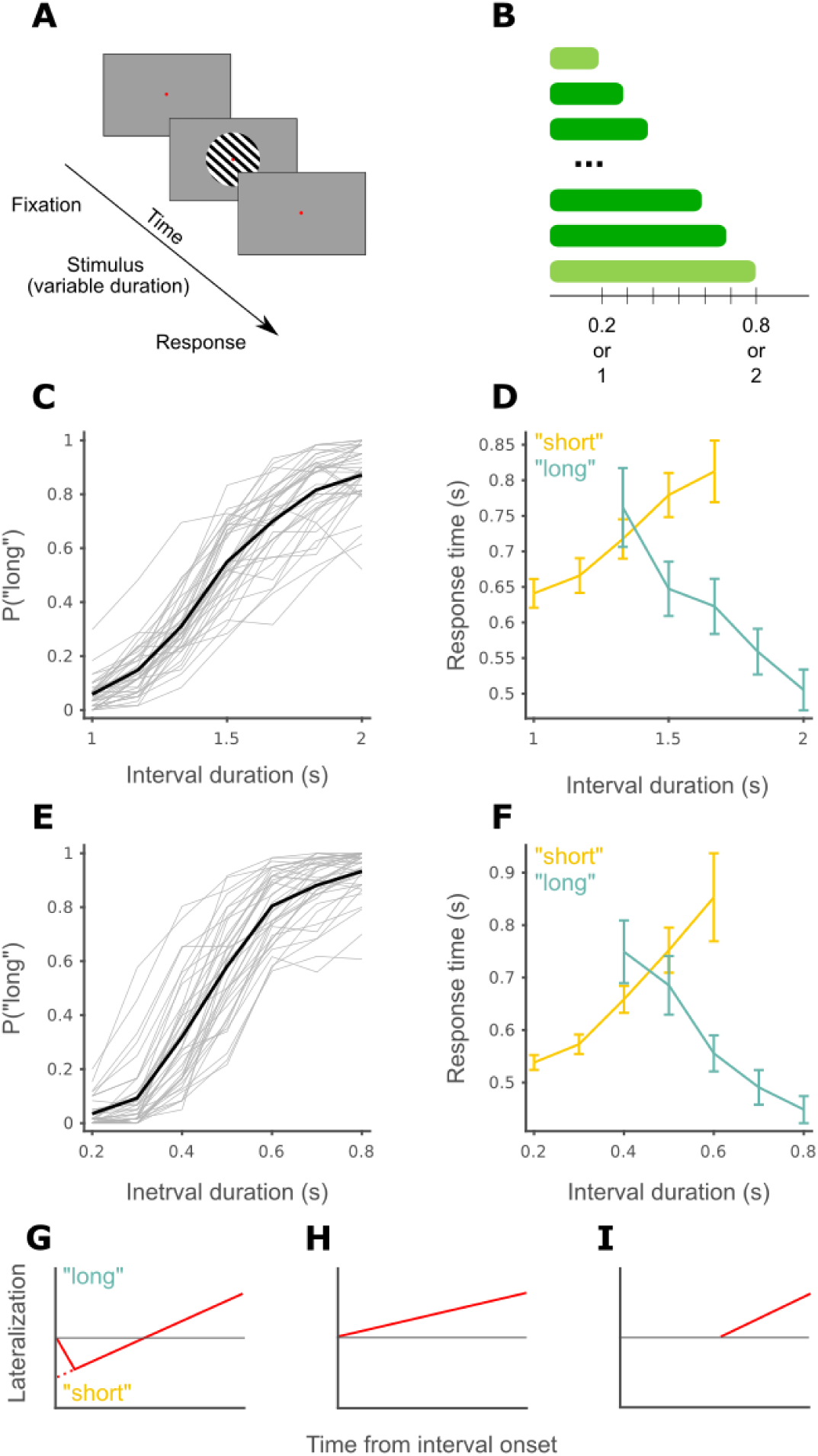
Temporal bisection task. **(A)** Schematic of a single trial. **(B)** Task design. The shortest and longest intervals, in light green, are the reference stimuli **(C)** Psychometric curve for the 1-2 s block. Grey lines depict the performance of single participants and thick black line is the group average. **(D)** Response time as a function of duration and response for the 1-2 s block. Group average and across participants SEM. Within each response, the two extreme intervals were not plotted as they contain very little “long” or “short” responses. **(E)** Same as **C** for the 0.2-0.8 s block. **(F)** Same as **D** for the 0.2-0.8 s block. **(G-I)** Hypothetical patterns of motor preparation following interval onset.

## Materials and Methods

This study reports an additional analysis for data that we first analyzed in a previous report (Ofir & Landau, 2022).

### Participants

Forty individuals (23 women, average age = 25 (SD 4.2)) participated in the experiment, corresponding to experiments 3a and 3b in the original study. Participants were recruited from the university community and were compensated for their time with either money (10 euro per hour) or class credit. All procedures were approved by the institutional review board of ethical conduct. Four participants did not complete one of the tasks (see Experimental Design and Statistical Analyses) due to technical reasons, 2 in each task, resulting in a dataset of 38 participants in each task.

### Stimuli and apparatus

Visual stimuli consisted of a square-wave grating presented in a circular window on a BenQ XL2420Z monitor running on 100 Hz (experiment 2) using Psychtoolbox (Kleiner et al., 2007) in Matlab (MathWorks, MA). The grating had a spatial frequency of 3 cycles per visual degree, a diameter of 8° visual degree and was positioned at the center of the screen. During the experiment, stimuli were presented for different durations (see experimental procedure) at two different levels of contrast, 100% and 50%. The gratings were presented randomly with a tilt of 45° or 135°, and a phase of 0°, 90° or 180°.

### Experimental Design and Statistical Analyses

The participants did a visual version of the temporal bisection task (**Figure 1A-B**). In this task, participants are first trained briefly to identify short and long reference intervals and are then requested to categorize intervals within that range as being more similar to the short or long references. The participants did the task twice, once with 0.2 s and 0.8 s as the short and long references, and once with 1 s and 2 s as the references, in two separate blocks. The test intervals were [0.2, 0.3, 0.4, 0.5, 0.6, 0.7, 0.8] and [1, 1.17, 1.33, 1.5, 1.67, 1.83, 2] seconds in each block respectively. The familiarization phase included 12 trials per duration. The test phase included a total of 420 trials, with 40 trials for each duration. During the test phase, a break was given to the participants every 84 trials (every ∼10 minutes). Each interval was presented 12 times within each block of 84 trials in random order. In the training phase, participants received feedback for each response, and within the test phase, participants received feedback only for their responses on the shortest and longest stimuli (the reference intervals). A red fixation point was displayed at the center of the screen (atop the gratings) throughout the entire experiment, excluding breaks. For a trial to start, participants had to fixate within a 1.5° radius of the fixation dot for a continuous second. Participants were asked to respond quickly and accurately after stimulus offset. Participants used a different hand to make “short” and “long” responses, allowing us to look at the dynamics of motor preparation as a window onto the cognitive processes underlying behavior in the task (O’Connell & Kelly, 2021). The response-hand mapping and block order were counterbalanced across participants.

### EEG acquisition

We recorded the EEG of the participants using a g.GAMMAcap (gTec, Austria) and a g.HIamp amplifier (gTec, Austria). The cap contained 62 active electrodes, positioned over the scalp according to the extended 10–20 system, with the addition of two active earlobe electrodes. We removed electrodes F9 and F10, as they often included strong muscle activity and were far from regions of interest. In addition, for all participants we recorded the horizontal electrooculogram (EOG) using passive electrodes placed at the outer canthi of both eyes and the vertical EOG using electrodes placed above and below the left eye. The EEG was continuously sampled at 512 Hz. We monitored the eye position using an infrared EyeLink camera (SR Research, Canada), sampling at 1000 Hz. The EyeLink signal and the EEG signal were time-aligned and stored for offline-analysis using a Simulink model (MathWorks, MA).

### EEG preprocessing

The EEG was referenced offline to the average of the earlobes. All offline preprocessing and analyses were done using a combination of FieldTrip (Oostenveld et al., 2011), EEGLAB (Delorme & Makeig, 2004) and custom Matlab code. Bad electrodes were removed by visual inspection. On average, 0.75% of electrodes per participant were removed (max 6.67%). None of the participants had bad electrodes within the region of interest (C3, C4, CP3, CP4). Slow drifts were removed using a spline-based approach (Ofir & Landau, 2022). Trials were defined from 500 msec before stimulus onset to 500 msec after the participant responded. Artifactual trials were removed using visual inspection guided by the summary statistics, implemented in ft_rejectvisual(). On average, 2.05% (max. 9.05%) and 2.45% (max. 6.43%) of trials were removed per participant in the short (0.2-0.8 s references) and long (1-2 s references) blocks, respectively.

### Motor signals analysis

The extraction of mu-beta lateralization closely followed the recent literature (Corbett et al., 2023). First, we applied a surface Laplacian transformation to the EEG data using the spherical splines approach (Perrin et al., 1989), implemented in ft_scalpcurrentdensity(). We identified the channels for each signal based on the grand average topographies (**Figure 2D**, **Figure 3E-F**).

**Figure 2:**
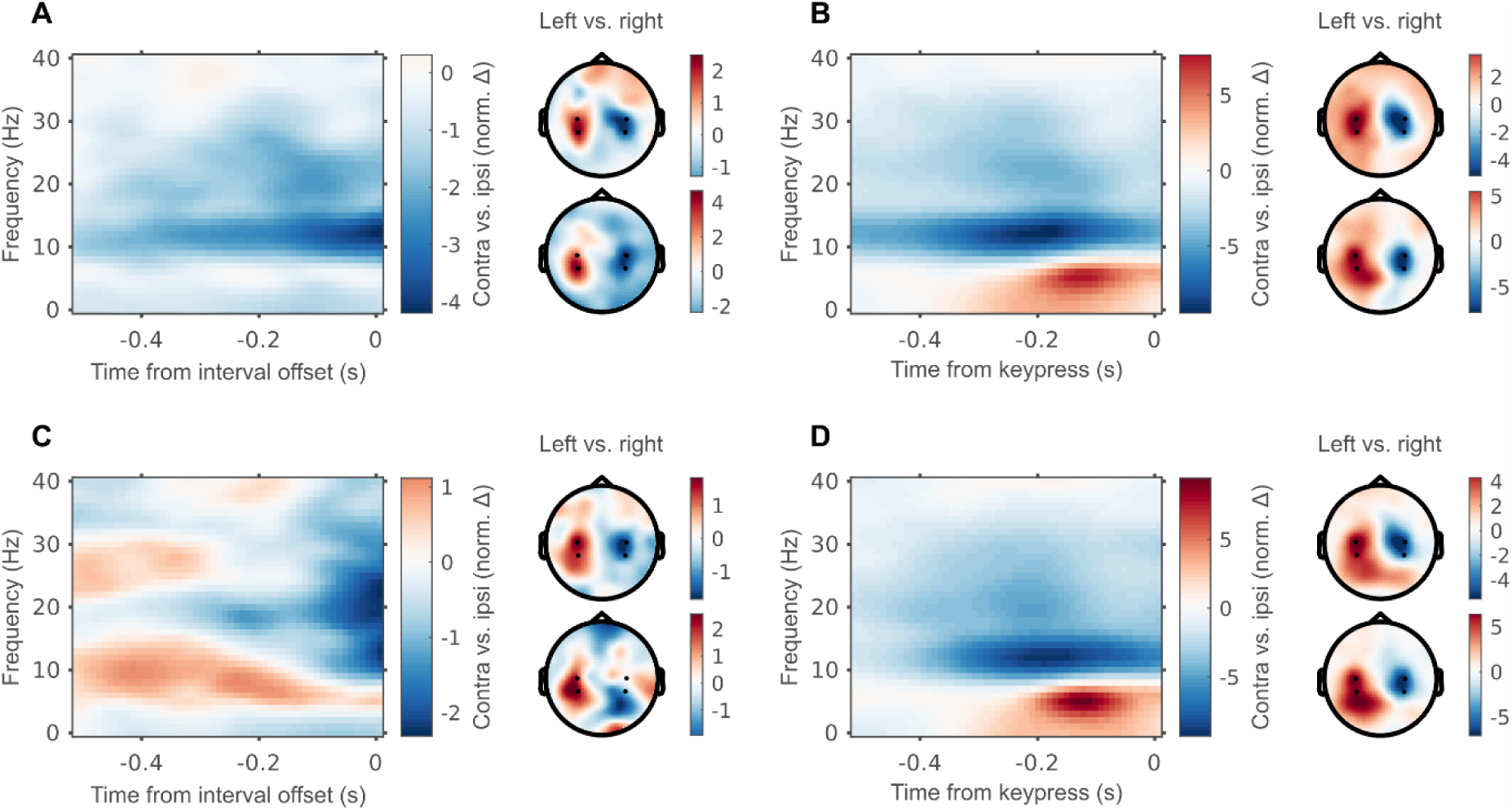
Mu-beta starts lateralizing before interval offset. **(A)** On the left, the lateralized time-frequency representation locked to interval offset for the 1-2 s block, using electrodes C3, CP3, C4 and CP4 (as described in the main text). On the right are the topographies of the alpha band (8-12 Hz, bottom) and beta band (14-30 Hz, top) amplitudes for left-hand vs. right-hand responses. **(B)** Same as **A**, but locked to keypress. **(C)** Same as **A**, for 0.2-0.8 s block. **(D)** Same as **B**, for 0.2-0.8 s block.

**Figure 3:**
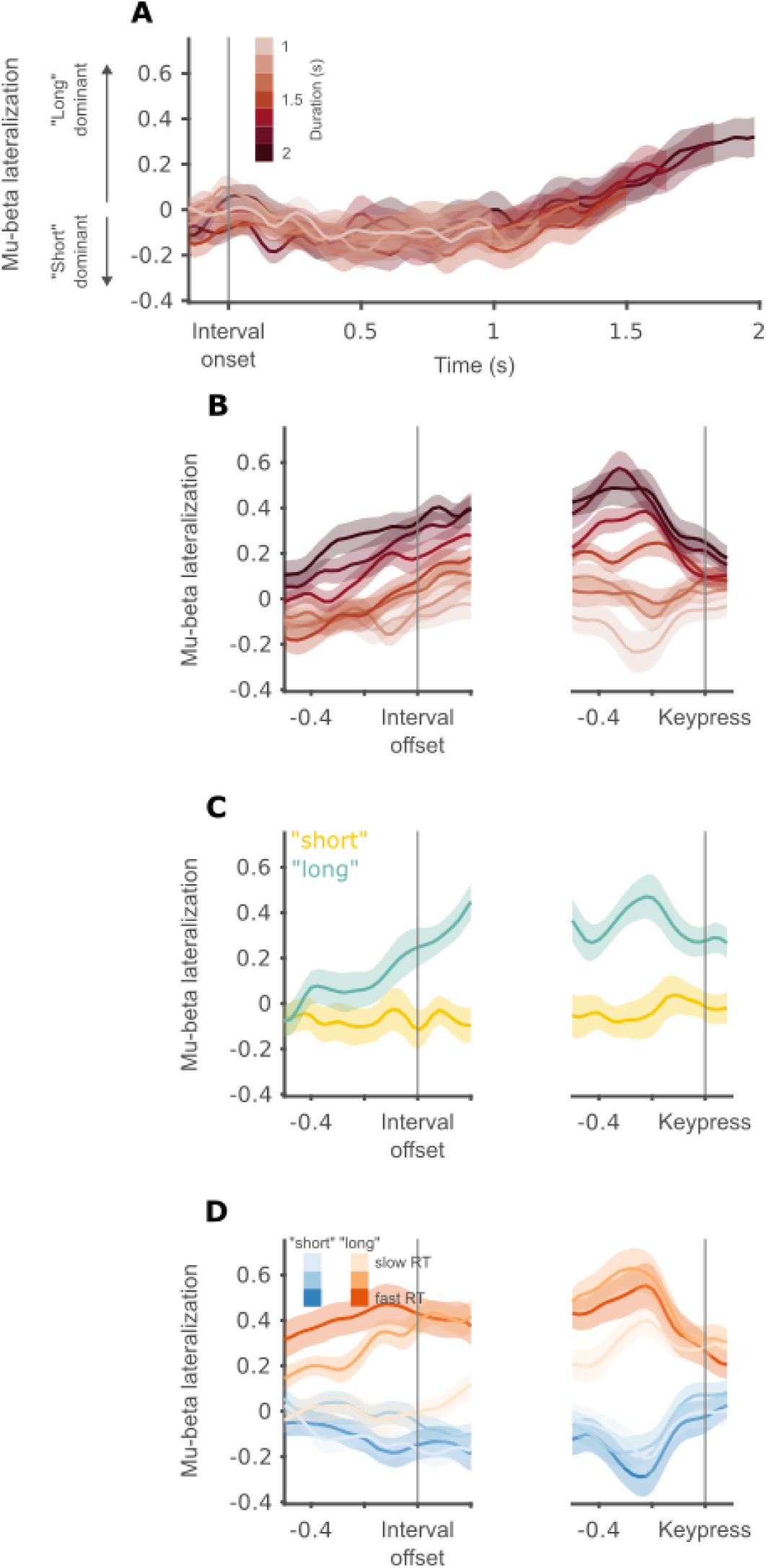
Mu-beta lateralization tracks “long” decisions, 1-2 s block. **(A)** Mu-beta lateralization traces as a function of interval duration, marked using color lightness. Grand average and between-participant SEM locked to interval onset. **(B)** Mu-beta lateralization traces as a function of interval duration, marked using color lightness. Left, grand average and within-participant SEM locked to interval offset. Right, grand average and within-participant SEM locked to keypress. **(C)** Mu-beta lateralization traces as a function of response (yellow for “short”, cyan for “long”) at individual bisection points. Subpanels as in **B**. **(D)** Mu-beta lateralization traces as a function response (blue for “short”, red for “long”) and RT (darker colors correspond to faster RTs). Subpanels as in **B**.

To validate that mu-beta is measurable as expected in our data, we computed TFRs before interval offset and keypress using a short-time Fourier transform, with 300 msec Hanning windows and 20 ms steps. We compared the spectrotemporal pattern of EEG between trials in which participants responded using their left hand, to those in which they used their right hand. In both blocks, of the 1-2 s intervals and the 0.2-0.8 s intervals, lateralization in the 8-30 Hz band peaked around 200 ms before keypress, at the expected channels (**Figure 2B and 2D**). Importantly, the spectrotemporal and spatial patterns of this lateralization were also evident before interval offset (**Figure 2A and 2C**). The offset-locked TFR in the 0.2-0.8 s block was not as clear, since in this case the intervals are much shorter, and the 0.5 s window extends beyond interval onset for many of the intervals we used. For the following analyses, we focused on mu-beta lateralization, defined as the difference between the 8-30 Hz amplitude in the channels contralateral to the “short” hand and in the channels contralateral to the “long” hand. That is, avg(C4, CP4) minus avg(C3, CP3) for participants who used their right hand for “long”, and avg(C3, CP3) minus avg(C4, CP4) for participants who used their left hand for “long”. This definition means positive values indicate lateralization towards the “long” hand, and negative values indicate lateralization towards the “short” hand. Using only the “alpha” (8-12 Hz) or “beta” band (14-30 Hz) yielded similar results (**Table 1&Table 2**). Given the temporal averaging inherent in frequency-domain analysis we used a window of 320-20 ms before keypress (a 300 ms window centered on 170 ms before keypress) and a window of -150-150 ms around interval offset to measure mu-beta lateralization. As we did not find significant lateralization in the pre-interval baseline period (One sample Wilcoxon signed-rank test, p = 0.180) we baselined the lateralization index.

**Table 1:** LMM results for the 1-2 s block. Each row contains results for a single model parameter. The table is divided into 2 parts, one for the offset window, and one for the keypress window. Each cell contains the estimate of the relevant beta coefficient and in parentheses the p value for a significance test against zero. Cells with significant coefficients (p < 0.05) are marked in bold.

|  | Offset |  |  | Keypress |  |  |
| --- | --- | --- | --- | --- | --- | --- |
|  | 8-12 Hz | 14-30 Hz | 8-30 Hz | 8-12 Hz | 14-30 Hz | 8-30 Hz |
| Intercept | **-0.25**<br>(0.045) | -0.02<br>(0.634) | -0.08<br>(0.210) | -0.20<br>(0.124) | **-0.17**<br>(**< 0.001**) | **-0.16**<br>(0.011) |
| Duration | -0.05<br>(0.673) | 0.02<br>(0.599) | 0.004<br>(0.939) | 0.14<br>(0.129) | 0.05<br>(0.222) | 0.08<br>(0.084) |
| Response | **0.67**<br>(**< 0.001**) | **0.31**<br>(**< 0.001**) | **0.37**<br>(**< 0.001**) | **1.02**<br>(**< 0.001**) | **0.69**<br>(**< 0.001**) | **0.68**<br>(**< 0.001**) |
| RT | -0.15<br>(0.367) | -0.07<br>(0.360) | -0.11<br>(0.207) | 0.11<br>(0.434) | -0.04<br>(0.504) | -0.01<br>(0.844) |
| Duration by response | 0.19<br>(0.242) | 0.04<br>(0.559) | 0.07<br>(0.386) | -0.14<br>(0.321) | -0.02<br>(0.763) | -0.06<br>(0.366) |
| RT by response | **-0.58**<br>(0.011) | **-0.28**<br>(0.004) | **-0.32**<br>(0.007) | **-0.59**<br>(0.003) | -0.08<br>(0.334) | **-0.20**<br>(0.042) |

**Table 2:** LMM results for the 0.2-0.8 s block. Each row contains results for a single model parameter. The table is divided into 2 parts, one for the offset window, and one for the keypress window. Each cell contains the estimate of the relevant beta coefficient and in parentheses the p value for a significance test against zero.

|  | Offset |  |  | Keypress |  |  |
| --- | --- | --- | --- | --- | --- | --- |
|  | 8-12 Hz | 14-30 Hz | 8-30 Hz | 8-12 Hz | 14-30 Hz | 8-30 Hz |
| Intercept | -0.155<br>(0.230) | -0.086<br>(0.127) | -0.108<br>(0.119) | **-0.262**<br>**(0.046)** | **-0.251**<br>**(<4e-6)** | **-0.229**<br>**(<7e-4)** |
| Duration | -0.146<br>(0.193) | -0.090<br>(0.058) | **-0.119**<br>**(0.033)** | 0.007<br>(0.945) | -0.036<br>(0.414) | -0.036<br>(0.486) |
| Response | 0.089<br>(0.411) | **0.247**<br>**(<8e-8)** | **0.201**<br>**(<3e-4)** | **0.868**<br>**(<6e-18)** | **0.800**<br>**(<2e-77)** | **0.737**<br>**(<4e-49)** |
| RT | 0.030<br>(0.851) | 0.100<br>(0.141) | 0.068<br>(0.395) | 0.207<br>(0.160) | 0.050<br>(0.426) | 0.087<br>(0.236) |
| Duration by response | **0.682**<br>**(<3e-05)** | **0.344**<br>**(<4e-7)** | **0.426**<br>**(<2e-7)** | **0.322**<br>**(0.029)** | 0.080<br>(0.204) | 0.141<br>(0.054) |
| RT by response | -0.260<br>(0.234) | **-0.285**<br>**(0.002)** | **-0.246**<br>**(0.024)** | -0.256<br>(0.205) | **-0.183**<br>**(0.033)** | **-0.197**<br>**(0.049)** |

The data visualizing the effect of decision (“short” vs. “long” trials for the same interval duration, **Figures 2F, 4D-E, 5D-E**) was calculated by taking the interval duration that was closest to the bisection point for each participant and extracting the signals of interest for those stimuli (Ofir & Landau, 2022). We averaged the signals within participants and then across participants for trials that were categorized as “short” and “long”, separately.

The data visualizing the combined effects of decision and RT (fast vs. slow responses, for “short” and “long” trials separately, **Figures 4F-G, 5F-G**) was calculated as follows: RTs were taken from all behavioral data (not removing trials with EEG artifacts), bar those with RT longer than 5 seconds (a cutoff we applied also for the EEG data, see Ofir & Landau, 2022). Within each response, we calculated 3 equal-size bins of RTs. We then averaged the signals within participants in each of the 6 pseudo-conditions (fast/medium/slow X “short”/”long”) and then across participants.

**Figure 4:**
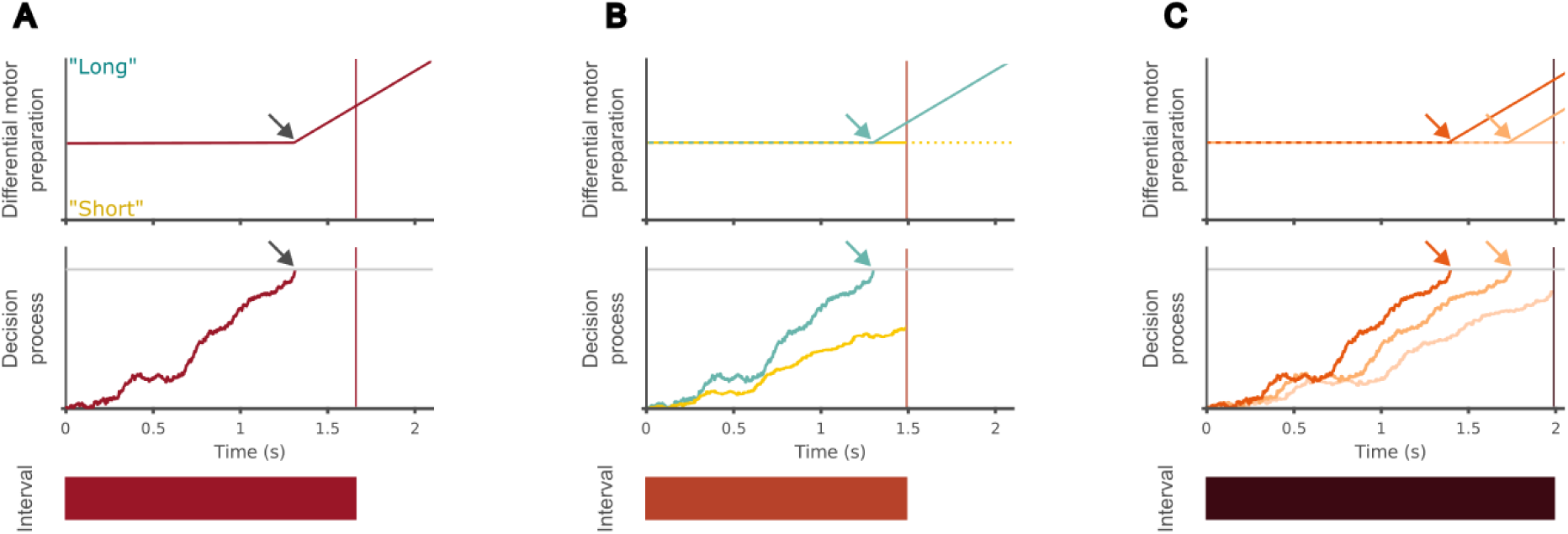
The two-stage model of temporal bisection. (**A)** Model description through an example trial. Red rectangle at the bottom depicts the time interval, lasting 1.67 s; the decision process is presented in the middle panel and includes a horizonal grey line signifying the decision boundary. For this trial, the accumulator reaches the boundary at 1.25 s after interval onset, and the interval is categorized as “long”. Motor preparation, in the top panel, is at zero until the boundary is reached, and then a “long” response is prepared. The vertical line in the decision and motor processes plots depict interval offset. The grey arrows mark the moment in which the accumulator reaches the decision boundary (in the middle panel) and the moment in which motor preparation begins (in the top panel). (**B)** Model prediction for “short” vs. “long” decisions, depicted in yellow and turquoise respectively, using two trials with an equal interval duration of 1.5 s. The accumulator reached the boundary in one trial (turquoise trace), so it is categorized as “long”. In the other trial we assume that the interval is categorized as “short” in the 2^nd^ stage, but we are agnostic about the dynamics of motor preparation in that stage. At interval offset, there is differential motor preparation towards the “long” hand in the “long” trial, but no differential motor preparation in the “short” trial. (**C)** Model prediction for RT effect in “long” trials. For example, a single interval, 2 s long, is presented 3 times. In one trial the boundary is reached before 1.5 s, in the second it is reached after 1.5 s and in the third the boundary isn’t reached at all. We assume that the third trial is categorized as “long” by the 2^nd^ stage. Reaching the boundary earlier means the motor process has more time to develop until interval offset, resulting in fasters RT and stronger lateralization at interval offset.

### Statistical tests

To test for significant lateralization during the first second of the interval in the 1-2 s block, we used a one-sample t-test against zero, with an FDR correction for multiple comparisons (Genovese et al., 2002).

To explore the factors affecting mu-beta lateralization, at interval offset and keypress separately for each block, we used linear mixed models implemented in Matlab’s fitlme(). All models, 4 in total, contained decision (categorical, “short” or “long” with “short” as the reference level), interval duration (continuous, scaled to vary between -1 and 1, with 0 the mean of range: 1.5 for the 1-2 s intervals, 0.5 for the 0.2-0.8 s intervals), and RT (centered within each participant and response category). We included random intercepts in all models. The model formula is:

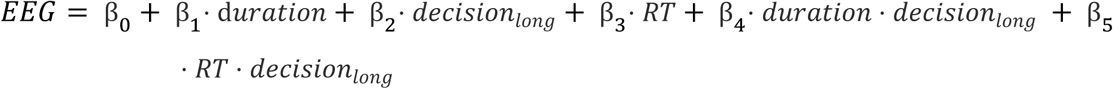

The model parameters, then, have the following interpretation:

1. β_0_ – lateralization associated with a “short” response at the mean duration and RT.
2. β_1_ – the effect of increasing interval duration from the mean to the longest for “short” responses.
3. β_2_ – the difference “long”- “short” decisions at the mean duration and RT.
4. β_3_ – the effect of increasing RT by 1 second for “short” responses.
5. β_4_ – the difference of increasing interval duration from the mean to the longest for “long” responses compared to the effect for “short” responses.
6. β_5_ – the difference of increasing RT by 1 second for “long” responses compared to the effect for “short” responses.

## Results

### Mu-beta lateralization tracks “long” but not “short” responses during the interval

We start by describing the dynamics of mu-beta lateralization in the 1-2 s bisection block. from interval onset and up to one second into the interval, mu-beta lateralization remained close to zero (no p_corrected_ < 0.05; **Figure 3A**). This finding rejects both the first scenario, predicting an initial lateralization towards “short” gradually switching to lateralization towards “long”, and the second scenario of a lateralization towards “long’ building from interval onset. The remaining scenario, the third we describe in the introduction, predicts lateralization will emerge after a “long” decision has been committed. After one second has elapsed from interval onset, a gradually increasing dominance of preparation to a “long” response is apparent. This increase can be visualized succinctly by aligning the traces to interval offset (**Figure 3B**). We next explore in more detail the factors underlying mu-beta lateralization in our task. As interval duration is strongly correlated with behavior in this task (**Figure 1C-F**), we used a mixed linear model to identify which experimental and behavioral variables explain mu-beta lateralization at interval offset: interval duration, participant’s response (“long”, “short”) and response time (RT), as well as the interaction terms of duration by response and RT by response (full statistical results are presented in **table 1&table 2**). We found that, once variability related to the behavioral output is considered, interval duration does not explain additional significant variability (duration predictor, β = - 0.08, p = 0.210; duration by response interaction, β = 0.07, p = 0.386). Behavioral output, on the other hand, displayed a rich pattern of correlations with mu-beta lateralization. First, there was significantly stronger lateralization towards the “long” hand for trials in which participants responded “long” compared to “short” (response predictor, β = 0.37, p < 0.001), but no significant lateralization for trials in which participants responded “short” (intercept, β = -0.08, p = 0.210). This effect is most clearly seen in trials with interval durations that are closest to individual bisection points (the interval for which participants respond “short” on approximately half of the trials). For a fixed duration, clear lateralization is seen before offset for trials with “long” responses. In contrast, for the same fixed intervals, no lateralization is found before interval offset for trials with “short” responses (**Figure 3C**). Beyond the strong effect of response, response time also predicted lateralization at interval offset. Importantly, the effect of response time was different for “short” and “long” trials. While for “short” trials we did not find a significant association between RTs and lateralization (RT predictor, β = -0.11, p = 0.207), for “long” trials faster RTs were associated with stronger lateralization at offset (RT by response interaction, β = -0.32, p = 0.007; **Figure 3D**).

We next tested whether duration, participants’ response and response time predict the level of lateralization prior to the keypress, using the same linear mixed model. The pattern at keypress differs somewhat from what we found at interval offset. As would be expected from a motor signal, we found significant lateralization for both “short” and “long” responses (intercept, β = -0.16, p = 0.011; response predictor, β = 0.68, p < 0.001; **Figure 4C**). Still, lateralization for “long” responses was larger in absolute terms when compared to “short” responses (paired samples Wilcoxon signed-rank test, p = 0.002). Lateralization was stronger for faster “long” responses, but there was no significance association between RTs and lateralization for “short” responses (RT predictor, β = -0.01, p = 0.844; RT response interaction predictor, β = -0.20, p = 0.042; **Figure 3D**). As at interval offset, interval duration did not significantly predict lateralization (duration predictor, β = 0.08, p = 0.084; duration response interaction, β = -0.06, p = 0.366; **Figure 3B**).

### Mu-beta lateralization reflects “long” decision boundary crossing

To summarize, mu-beta lateralization during the timed interval presents two important features. First, it starts neutral, and remains so during the first second of the timed interval, which corresponds to the shortest interval in the set. From that point and towards the interval’s offset, lateralization gradually increases towards “long”. Second, the level of lateralization at interval offset is associated with the RT of “long” responses. The first result rules out the first 2 scenarios we described in the introduction. Namely, that lateralization will be biased initially towards the “short” hand or that it will start neutral and immediately grow towards the “long” hand. Instead, the results are in line with the scenario that lateralization only starts after a decision has been made. We now turn to explain our results within the framework of the two-stage decision process that was hypothesized to underlie psychophysical performance in temporal bisection (Figure 4A; Balcı & Simen, 2014). In the first stage of this model, a noisy accumulator starts with interval onset, and runs until either it reaches a decision boundary or the interval ends. If the boundary is reached, the interval is categorized as “long” and preparation for the suitable motor response initiates. If it does not, a second stage starts at interval offset. In this stage, the value of the accumulator at interval offset is compared to an internal representation of a threshold between the reference durations. This second stage has two bounds, one for each response (“short” and “long”), and is thought to reflect resampling from memory, as the evidence itself (i.e. interval duration) is no longer directly available (Shadlen & Shohamy, 2016; van Ede & Nobre, 2024).

The basic, and distinguishing, assumption of the two-stage model is that an interval can be categorized as “long” before it ends (i.e., as soon as the decision boundary of the 1^st^ stage is reached) but can only be categorized as “short” after it ends. This is supported by our finding that “long” responses, but not “short” responses, are associated with significant lateralization at interval offset (**Figure 4B**). Furthermore, due to the inherent variability of the accumulator during the 1^st^ stage, there will be trial-to-trial variability in the time it takes to reach the boundary. In trials in which the accumulator rises quickly and reaches the boundary early, motor preparation can start earlier. This will translate into stronger lateralization at interval offset and faster responses. In other trials, the boundary will be reached later, but still before interval offset. This will translate into somewhat weaker lateralization at interval offset, and slower RTs. Finally, in some trials, the accumulator will not reach the boundary at all, but the interval will still be categorized as “long” by the 2^nd^ stage. In those trials we expect no lateralization at interval offset, and the slowest RTs (**Figure 4C**). This pattern was indeed what we found: strongest lateralization for the faster “long” responses, and essentially no lateralization for the slowest “long” responses (**Figure 3D**).

If the mu-beta pattern we find indeed reflects decision boundary crossings, it should scale with the position of the boundary. The position of the boundary itself shifts with the reference intervals, or in other words, with the temporal context. To test this, we turned to the second block in the experiment, in which the same participants performed the same temporal bisection task on intervals lasting between 200 and 800 ms. The patterns that emerge match those we found using longer intervals but scaled in time to fit the much shorter durations. We used the same statistical model on the data of this block, and the results generally replicate those we found in the 1-2 s block. At interval offset, “long” but not “short” responses were preceded by significant lateralization (intercept, β = -0.11, p = 0.119; response predictor, β = 0.20, p < 0.001; **Figure 5B**). Lateralization was significantly stronger for faster “long” responses, while we did not find a significant association between RT and lateralization in “short” trials (RT response interaction predictor, β = -0.25, p = 0.024; RT predictor, β = 0.07, p = 0.395; **Figure 5C**). The results of the 0.2-0.8 s block diverge from the 1-2 s block with respect to the effect of duration on lateralization. In the 1-2 s block we did not find a significant effect of interval duration. Here we found, somewhat counter-intuitively, that longer intervals were associated with stronger lateralization towards the “short” side in “short” trials (duration predictor, β = -0.12, p = 0.033), and with stronger lateralization towards the “long” side in “long” trials (duration response interaction, β = 0.43, p < 0.001; **Figure 5A**). We suspect this is an artifact, as in this experiment the peri-offset window is long relative to the intervals and extends back in time close to the onset for the shorter intervals, in which “short” responses are more prevalent.

**Figure 5:**
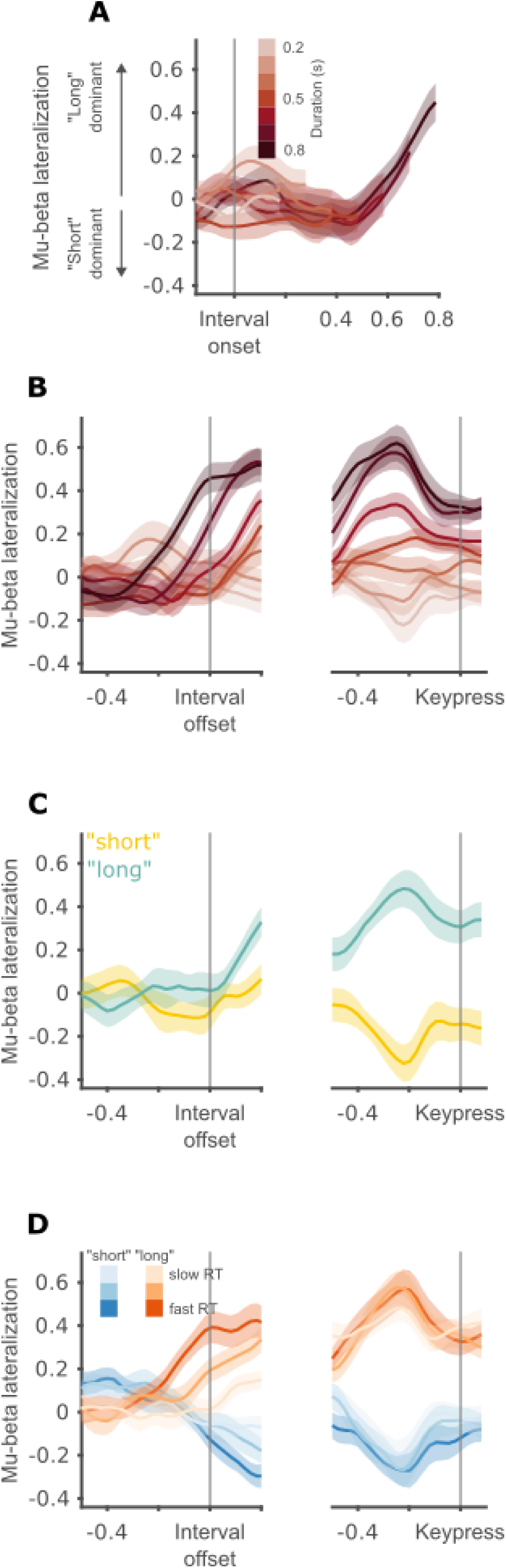
Mu-beta lateralization tracks “long” decisions, 0.2-0.8 s block. **(A)** Mu-beta lateralization traces as a function of interval duration, marked using color lightness. Grand average and between-participant SEM locked to interval onset. **(B)** Mu-beta lateralization traces as a function of interval duration, marked using color lightness. Left, grand average and within-participant SEM locked to interval offset. Right, grand average and within-participant SEM locked to keypress. **(C)** Mu-beta lateralization traces as a function of response (yellow for “short”, cyan for “long”) at individual bisection points. Subpanels as in **B**. **(D)** Mu-beta lateralization traces as a function response (blue for “short”, red for “long”) and RT (darker colors correspond to faster RTs). Subpanels as in **B**.

Before keypress, our statistical analysis provided results very similar to those seen in the 1-2 s block. We found significant lateralization for “long” as well as “short” responses (intercept, β = -0.23, p < 0.001; response predictor, β = 0.74, p < 0.001; **Figure 5B**). Lateralization before keypress was stronger for faster “long” responses, while lateralization was not significantly associated with RT for “short” responses (RT response interaction predictor, β = -0.20, p = 0.049; RT predictor, β = 0.09, p = 0.236; **Figure 5C**). Interval duration did not significantly predict lateralization (duration predictor, β = -0.04, p = 0.486; duration response interaction, β = 0.14, p = 0.054).

## Discussion

Timing is a basic cognitive ability which underlies essentially all behavior. We studied how duration is internally represented using the temporal bisection task. This task involves several neural processes, which enables exploring the neural mechanisms of time perception from multiple perspectives. To explore the dynamics of temporal decision formation we used mu-beta lateralization, which reflects relative motor preparation (O’Connell & Kelly, 2021; Pfurtscheller & Lopes Da Silva, 1999).

From interval onset and up to at least the shortest interval has elapsed, mu-beta was not lateralized towards any choice. This contrasts with two possible scenarios in which elapsed duration is tightly linked to motor preparation, predicting either an initial preparation to “short” responses (Leon & Shadlen, 2003) or a slope towards “long” responses that starts immediately after interval onset. The delayed buildup we report is consistent with the third scenario, in which motor preparation starts only after “long” decisions are being made. This contrasts with multiple studies which found motor preparation reflects accumulated evidence and not only final decision (Afacan-Seref et al., 2018; Balsdon et al., 2023; Steinemann et al., 2018; Wilming et al., 2020; Wyart et al., 2012). A possible explanation is that elapsed duration is not considered by the brain as “evidence” in the same way many other aspects of stimuli are, such as contrast or motion direction (O’Connell et al., 2012). In other words, temporal decisions might not be implemented in the brain simply by plugging the output of an internal clock into the mechanisms of perceptual decision-making (McCone et al., 2023).

Furthermore, our data reveal properties of the dynamics of temporal decisions. First, we provide here direct neural evidence for the ubiquity of “pre-committals”, that is “long” decisions made before interval offset. This is evident from the fact that “long” decisions display significant lateralization even before the mean interval has elapsed (**Figure 4C**). While this property may sound trivial, models of temporal bisection typically assume that decisions are only made at interval offset (Gibbon, 1981; Kopec & Brody, 2010; Machado et al., 2009). Even in the bounded accumulation model that does consider the existence of such decisions, and that we take here as our main theoretical reference, pre-committals are thought to be relatively rare (Balcı & Simen, 2014). Indeed, the common method of fitting a separate DDM for each duration implicitly assumes that pre-committals are negligible, as RTs for pre-committals will only reflect motor time, and not any bounded accumulation process. The strong association of RTs and mu-beta lateralization at interval offset is further evidence for the link between committing to a “long” decision and motor preparation.

We found that absolute lateralization before keypress was stronger for “long” compared to “short” responses. A possible explanation is that lateralization before keypress is greater for responses that can be planned in advance (Twomey et al., 2016). This can be tested by delaying responses using a response cue. Delaying responses might also provide a cleaner view on the second stage of decision making, after interval offset.

Despite being a commonly studied signature, the exact role of the mu-beta band is unknown. It is debated whether and how the mu rhythm, with its strong harmonic in the same band as the beta rhythm, and the beta rhythm can be non-invasively distinguished (Rodriguez-Larios & Haegens, 2023; Schaworonkow, 2023; but see Cheyne, 2013). Studies rarely report both “alpha” (8-12 Hz) and “beta” (13-30 Hz) bands separately, but when they do, activity in both bands is highly similar (Boettcher et al., 2021; Rogge et al., 2022). This similarity suggests that amplitude changes in the beta band in the type of decision-making tasks described here are a result of dynamics of the mu rhythm, and not a separate contribution of beta rhythms. Understanding which cognitive and neural processes are related to mu versus beta rhythms is a critical step in ultimately elucidating their computational roles.

In summary, our data highlights the importance of a central characteristic of bounded accumulation models of timing: that temporal decision making is dynamic (Balcı & Simen, 2024). Beyond explaining an additional source of RT variability in temporal bisection, we believe this characteristic is critical when examining neural activity in timing tasks. Additionally, our data also show that while behavior in temporal bisection can be adequately fitted by decision making models, the neural implementation is not identical to that found in many decision-making studies. Considering timing behavior as resulting from an orchestration of multiple dynamical processes is a fruitful framework for exploring its neural mechanisms, and is line with current views of other cognitive capabilities (e.g., selective attention, Nobre & Van Ede, 2023). In the temporal bisection task, a motor preparation process tracks “long” decisions made during the interval. After the interval ends, an additional evidence accumulation process reflects the formation of “short” decisions (Ofir & Landau, 2022). This second process hypothetically represents sampling from memory, as the interval is already over and no additional evidence is present in the environment (van Ede & Nobre, 2024). The next clearest goal in our view is finding an online signature of elapsed duration, a process that presumably starts at interval onset and ends when mu-beta lateralization begins.

## Acknowledgements

The authors would like to thank Noa Itzhaki, Yoel Gordon and Gal Samuel for assistance in data acquisition. We thank the members of the Brain Attention and Time Lab (PI: A.N.L) and the members of the Dynamic Cognition research group (PI: Dr. Assaf Breska) for their input on the work. The Brain Attention and Time Lab (PI: A.N.L.) is supported by the James McDonnell Scholar Award in Understanding Human Cognition, ISF grant 958/16. This project has received funding from the European Research Council (ERC) under the European Union’s Horizon 2020 research and innovation programme (grant agreement no. 852387).

